# “*Punarnavayolepa Choornam*” in Iron Deficiency Anaemia management: Pharmaceutic insights and biological activity

**DOI:** 10.1101/2025.10.08.681069

**Authors:** Jyothish Gopinathan Nair, TS Rajani, Deepthy Mohan, Puthanpurakal Indu Chandrabose, Varsha Wilson, Atheena Biju Shyni, Remya Venugopala Menon, Ashi Augustine, Yadu Narayanan Mooss, Subramanya Kumar Kukkupuni, Sheela Karalam Balakrishnan, Chethala N Vishnuprasad, Asish Gopinathan Remanikutty

**Affiliations:** Vaidyaratnam Ayurveda Research Institute, Thaikkattussery, Thrissur, Kerala 680306; Ayurveda Biology and Holistic Nutrition, The University of Transdisciplinary Health Sciences and Technology (TDU), No.74/2, Jarakabande Kaval, Post: Attur, Via Yelahanka, Bengaluru - 560 106, Karnataka, India; Department of Botany, Christ College (Autonomous), Irinjalakuda, Thrissur, Kerala, 680125; National Institute of Advanced Studies (NIAS), Indian Institute of Science Campus, Bengaluru - 560012, Karnataka, India

## Abstract

**Background and Aim:** Indigenous medical systems employ unique pharmaceutical techniques to meet the therapeutic needs. In the Indian System of Medicine, *Ayurveda*, a unique method called “*Ayolepam*” is described that facilitates efficient iron absorption from its source to the formulation. *Punarnavayolepa Choornam* (PC), prepared from *Boerhavia diffusa* L., is one such novel *Ayolepam* formulation developed by a traditional *Ayurveda* school of South India, widely used in the clinical management of iron deficiency anaemia. This study aims at scientifically evaluating the *Ayolepam* technique by assessing the iron binding and bioavailability properties of PC, prepared from the leaves, root and whole plant of *B. diffusa*, using *in-vitro* model systems..

**Experimental Procedure:** The iron content in both raw and processed samples of PC was quantified using Inductively Coupled Plasma Mass Spectrometry (ICP-MS). A simulated *in-vitro* digestion model employed to assess the release of bioavailable iron from the formulation. Subsequently, iron bioavailability was evaluated using the Caco-2 cell model of human intestinal epithelium following the ferrozine method.

**Results and Conclusion:** PC preparation from leaves, whole plant and roots of *B. diffusa*, showed significant increase in iron content compared to the raw material. *In-vitro* digestion studies confirmed the efficient release of bioavailable iron from these formulations, and subsequent Caco-2 cell assays confirmed their iron bioavailability properties. In conclusion, these findings provide a preliminary scientific basis for this novel and unique pharmaceutical design from a traditional school of *Ayurveda*, supporting its clinical application.

## 1. Introduction

Preparation of a formulation satisfying all pharmaceutic principles is extremely important and sometimes even challenging [1]. Before the discovery of modern techniques and instruments, traditional medical practices across the globe established their own unique and innovative methods for preparing formulations to meet the pharmacodynamic and pharmacokinetic requirements [2]. *Ayurveda*, the Indian System of Medicine (ISM), has described numerous methods to prepare drug formulations with optimal safety and efficacy [3] [4]. One such method of drug preparation is called “*Ayolepam*”, wherein the raw materials (or ingredients) are ground into a paste and smeared over a cast iron sheet and dried. The powder is scrapped off and used for clinical administration [5]. The word *Ayolepam* is derived from two Sanskrit words ‘*Ayo*’ (iron) and ‘*Lepam*’ (paste, ointment, smearing on a surface). *Ayurveda* describes the iron used in *Ayolepam* procedure as *‘Loha*’ [6]. Literature references like *Rasa Tarangini* and *Rasa Ratna Sammucchaya*, two classical texts on pharmacology and pharmaceutics of metals and minerals, mentions three types of *Loha* like *Munda Loha, Teekshna Loha*, and *Kaanta Loha [7] [8] [9]*. While in modern chemistry parlance these are similar, *Ayurveda* considers *Kaanta Loha* to be the most therapeutically potent type followed by *Teekshna Loha* and *Munda Loha*. Nevertheless, current practice uses *Teekshna Loha* (the mild steel iron) most commonly due to its easy availability [10].

Kerala, the southern state of India, is known for its unique schools of *Ayurveda* practice with several unique formulations and procedures for disease management [11]. One such unique school is called the “*Ashta Vaidya*” tradition, where the professionals trained in this school are known to have expertise in all eight branches of *Ayurveda* viz. Internal Medicine (*Kaya Chikitsa*), Surgery (*Shalya Tantra*), ENT and Ophthalmology (*Shalakya Tantra*), Pediatrics and Obstetrics (*Kaumarbhritya*), Toxicology (*Agada Tantra*), Psychiatry (*Graha Chikitsa*), Rejuvenation and Geriatrics (*Rasayana Chikitsa*), and Aphrodisiac therapy (*Vajikarana*) [12] [13] [14]. *Punarnavayolepa Choorna* (PC), the formulation of focus in this study, is one of the unique preparations from a renowned *Ashta Vaidya* school in Kerala, prepared following the pharmaceutic principles of *Ayolepam* technique. PC is a single-herb based preparation using the roots of *Boerhavia diffusa* Linn. (Family: Nyctaginaceae), referred to as *Punarnava* in *Ayurveda*. It is a well-known medicinal herb in ISMs and is also used in South America and Africa. Different parts of this plant, primarily the roots, are used for managing gynaecological diseases, hepatoprotection, and gastrointestinal disorders. *Boerhavia diffusa* Linn. is an important herb in *Ayurveda*, recognized as a *Rasayana*, and is found to be a key ingredient in 35 formulations. [15] [16]

The most common indication of PC is in the management of *Pandu Roga*, a disease described in *Ayurveda* whose clinical manifestations are comparable to the clinical manifestations of Iron Deficiency Anaemia (IDA). A notable pallor or yellowish-white colouring of the skin, signifying a decreased haemoglobin level and/or a drop in the number of red blood cells (RBCs), is characteristic of both *Pandu Roga* and IDA [17]. Globally, iron deficiency is recognised as the cause of 50% of anemia cases. It ranks 9th among the 26 risk factors in GBD 2000, contributing to over 841,000 deaths and 35 million disability-adjusted life years (DALYs) lost-primarily affecting Africa and Asia, which is 70% of global burden [18]. The most recommended strategies for managing IDA are dietary changes and iron supplements [19]. As diet is the only source of iron for the human body, and the bioavailability of iron from diet is too low due to various reasons, plus impairments in iron bioavailability leading to diseases like IDA, intervention of iron-folic acid like supplements are becoming imperative. However, problems like poor absorption, gastrointestinal side effects and low patient compliance frequently limit the use of oral iron supplements [20] These challenges corroborate the need of better, safe and culturally acceptable iron bioavailability enhancers for masses [21]. PC, a unique preparation of *Ashta Vaidya* tradition of *Ayurveda* is a choice in this direction, which is administered with buttermilk as an adjuvant. Although PC has been used for several years for the clinical management of IDA-like disease manifestations, there are no systematically documented data available on its clinical efficacy and mode of action.

In this background, the present study attempts to examine the pharmaceutic and pharmacological properties of PC emphasizing on its ability to enhance iron bioavailability in Caco-2 cells, a model of human intestinal absorption. Our study uses an in-house prepared PC using the roots, leaves and the whole plant of *Boerhavia diffusa* Linn. The formulation is analysed for its Physicochemical Analysis, Nutritional profiling, and iron bioavailability using various models.

## 2. Materials and Methods

### 2.1 Plant material, cell line and fine reagents

The *B. diffusa* plants were taxonomically identified by botanists of *Vaidyaratnam Ayurveda Research Institute* (VARI) and the leaves, roots and the whole plant were collected separately from the raw material storage section of Vaidyaratnam Ousadhasala, Thaikkattuserry, Ollur, Thrissur, Kerala. Caco-2 (human colorectal adenocarcinoma cell line) was a kind gift from Dr. Raghu Pullakhandam, National Institute of Nutrition, Hyderabad. Minimum Essential Medium (MEM - Cat. No. 11095072) and Foetal Bovine Serum (FBS - Cat. No. 10270106) were purchased from Thermo Fisher Scientific Inc. All fine chemicals for various bioassays were purchased from Sigma-Aldrich (St. Louis, MI).

### 2.2 Mild Steel Iron Trays

The standard mild-steel iron sheets were procured from a commercial vendor (M/s. Shankara Build Pro, Thrissur) and the in-house maintenance team of *Vaidyaratnam Oushadhashala*, Ollur, Thrissur, Kerala, India manufactured the sheets into trays with 40cm

× 40cm × 6 cm length × width × height) were used.

### 2.3 *Punarnavayolepa Choornnam* preparation

The plant parts collected were washed thoroughly with clean water to remove any debris present. The roots were dried and powdered, and made into a paste with a sufficient amount of water. The leaves and the whole plant were made into a paste without drying. All three samples (Leaves, Root and Whole plant) were smeared evenly on three separate iron trays and were dried in an oven at 40°C. The complete drying time for root, whole plant and leaf *Ayolepam* preparations were 66 hrs, 113 hrs and 88 hrs respectively. The powders were labelled and stored in air-tight glass containers for further studies.

### 2.4 Pharmacognosy Studies

Both raw materials and *Ayolepa* preparations were subjected to pharmacognosy analysis. Macroscopic characteristics of plants were documented through visual examination of leaf, stem and root. Transverse slices (T.S.) of the leaf, stem, and root were prepared for histological investigations in order to examine anatomical structure. To find distinctive microscopic characteristics, powdered samples of the leaf, stem, and root were subjected to powder microscopy. Powder microscopy was also used on Ayolepam formulations made from the three parts of the plant.

### 2.5 Physicochemical Analysis

Total Ash and Loss on Drying (LOD) were calculated using the methods described in the Ayurvedic Pharmacopoeia of India [22]. The Anthrone technique, as outlined in Sadasivam, S. and Manickam (2008) A Biochemical Method. 3rd Edition, New Age International Publishers, New Delhi. was used to measure total carbohydrates [23].

### 2.6 Micro and Macro Nutrient profiling of the Samples

For the nutrient profiling, 2gms of the samples were weighed and heated in a muffle furnace at 450°C for two and a half hours to make ashes of the sample. This ash is then treated with 5mL of 2N HCl, and slowly heated to ensure complete solubilization. The solution is left overnight undisturbed for complete reaction, and any insoluble residues were removed by filtration using filter paper. The filtered solution is transferred to a 50 ml volumetric flask, and made up to 50mL using distilled water. Standard solutions of calcium chloride, sodium chloride, and potassium chloride at linear concentrations of 25 ppm, 50 ppm, and 100 ppm are used to calibrate a flame photometer for analysis. To provide accurate and trustworthy nutritional evaluation, the prepared sample solution is lastly examined to ascertain its mineral content.

### 2.7 Simulated *in-vitro* digestion of samples

The *in vitro* digestion of PC was performed following the protocol standardized in the lab with minor modifications to suit the samples [24] The electrolyte solutions for simulated gastric fluid (SGF) and simulated intestinal fluid (SIF) were prepared as reported by Butala [24]. The three varieties of PC formulations (prepared as mentioned in section 2.3) were taken (0.5gm of each preparation) in 50 mL falcon tube and suspended in 12.5 mL of SGF supplemented with 2500 U/mL pepsin and 0.16 mM CaCl_2_.2H_2_O and mixed thoroughly. The pH was adjusted to 2.0 by adding 6N HCl and the tubes were incubated at 37°C for 2 hrs in a shaking water bath. Following gastric digestion, the gastric chyme was mixed with 12.5mL SIF containing pancreatin-bile solution (final concentration of 500 μg/mL pancreatin and 3 mg/mL bile) and 0.6 mM CaCl_2_.2H_2_O. The pH was adjusted to 7.0 by adding the required amount of 5N NaOH and incubated at 37°C in a shaking water bath for another 2 hrs. After complete digestion, the samples were heat inactivated by keeping them at 65°C water bath for 30 minutes to stop all enzymatic activities. The digests were centrifuged at 5000 rpm for 15 mins and filtered and the supernatant was collected and stored at −80°C for further use.

### 2.8 Iron Estimation Using ICP-MS

For the determination of iron by ICP-MS (Agilent), samples (minimum 1.2 mL in vial) are introduced via a concentric nebulizer with peristaltic uptake (~0.1–0.2 mL/min) under standard plasma conditions (RF power 1550–1600 W, sampling depth 8–10 mm, carrier gas 1.0–1.1 L/min, makeup gas ~0.1 L/min). Helium collision mode (He KED, 4–5 mL/min, 3–5 V) is used to remove ArO^+^ interference, with Fe monitored at m/z 56 (primary) and m/z 57 (confirmation). Rh-103 (or Sc-45/Ge-72) is employed as the internal standard. Dwell time is ~0.1 s with 30–50 sweeps and triplicate replicates per isotope. Calibration is prepared in 1% HNO_3_ over an appropriate range (e.g., 1–100 µg/L) with IS added, and blanks, continuing calibration verification, and independent check standards are analyzed at regular intervals. Rinse solution is 1–2% HNO_3_ between samples, with extended rinse after high concentration runs. Results are reported directly from software with dilution factor corrections applied [25].

### 2.9 Evaluating the iron bioavailability by Caco-2 cells following the ferrozine method

The Caco-2 cells were maintained in MEM media containing 10% FBS and essential amino acids. For the assays, the cells were cultured in 12-well plates for 21 days by changing the media every 48 hr. On the day of assay, the cells were treated with 500uL of growth media containing 150µg/mL of FeSO_4_ along with varying concentrations (30%, 20% and 10% v/v) of the *Ayolepam* digests. Cells treated with 150µg/mL of FeSO_4_ and 500µM ascorbic acid was used as experimental control (positive) and cells treated with respective concentrations of *Ayolepam* digests without FeSO_4_ was used as a control to see the iron bioavailability directly from the sample. The cells were incubated at 37°C for 18 hrs. After the incubation, the media is discarded and the cells were rinsed with 500 µL of ice-cold saline solution (140 mM NaCl, 0.9%) to remove residual media. After washing, 500 µL of stop solution (140 mM NaCl containing 10 mM PIPES) was added to each well and incubated for 1–2 minutes to stop the reaction. Followed by this, the cells were washed twice with 400µL of removal solution (stop solution + 5 mM bathophenanthroline disulfonic acid). After the final wash, 150 µL of solubilization solution (0.5 N NaOH) was added to each well, and the cells were completely scraped and transferred into microcentrifuge tubes. For complete lysis and solubilization, the cells were incubated at 60°C in a water bath for 30 minutes. Total cell protein of the different samples was estimated using the Bradford assay.

For estimating the intracellular iron in different groups, 200µL of freshly prepared Iron releasing solution (a 1:1 mixture of 1.4 M HCl and 4.5% KMnO_4_) was added to the lysates, followed by incubation for 2 hours at 60°C in a water bath. After incubation, 100µL of, iron detection reagent (6.5 mM ferrozine, 2.5 mM ammonium acetate, and 1 M ascorbic acid) was added to each tube and the reaction mixtures were mixed thoroughly and centrifuged at 4000 rpm for 5 minutes.100 µL of the supernatant was transferred to to 96-well plate, in duplicates, and the absorbance was measured at 550 nm using a UV-visible spectrophotometer to determine iron concentration. An iron standard curve was prepared from serially diluted 1 mg/mL FeSO_4_ and the total iron absorbed by the cells was represented as ‘µg of FeSO_4_ Eq iron/mg of protein’. [26] [27]

## Results

### 3.1 Pharmacognosy Studies

#### 3.1.1 Morphology of B. diffusa

The leaves are grouped in opposing, unequal pairs, featuring a rounded or slightly pointed tip. The upper surface is green and glabrous, whereas the underside is pale, occasionally exhibiting a pinkish tint on the dorsal side. The leaf margins are entire or slightly undulate. The stem is slender, cylindrical, and stiff, exhibiting a greenish-purple coloration and swelling at the nodes, with a minutely pubescent texture. The root system is well-developed, fairly long, and somewhat tortuous, with a cylindrical shape measuring between 0.2 and 1.5 cm in diameter. The roots exhibit a yellowish-brown to brown coloration. The plant produces tiny, pink blooms that are either short-stemmed or almost sessile. The upper portion of the flower appears pink and funnel-shaped, and the inflorescence is made up of terminal and axillary panicles grouped in a corymb pattern. The fruit has a structure that is bluntly five-ribbed. These physical characteristics aid in the plant’s identification and pharmacognostic assessment. (Figure 1)

**Figure 1.**
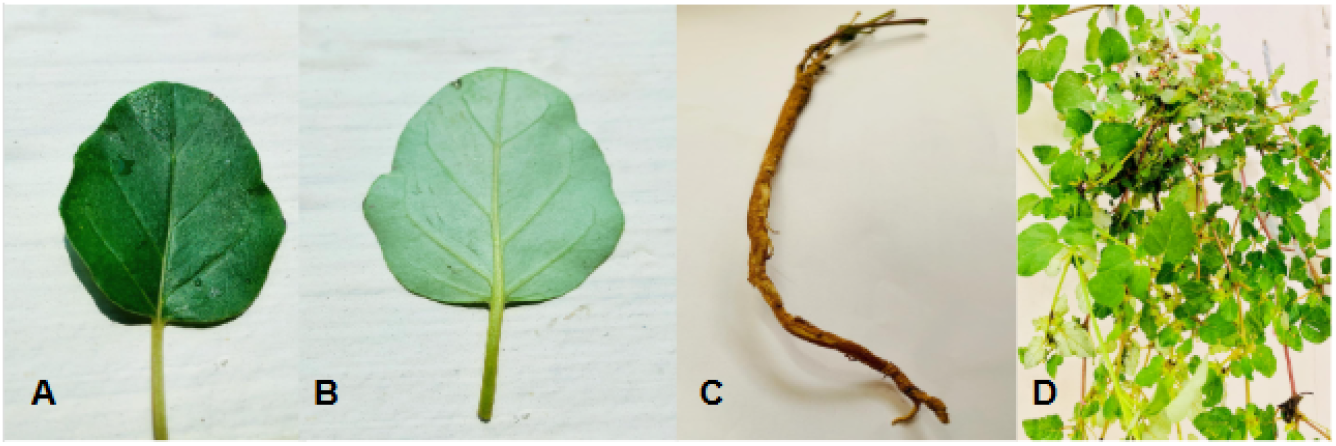
Morphological characteristics of the plant. **(A)** Dorsal (adaxial) surface of a representative leaf. **(B)** Ventral (abaxial) surface of the leaf, highlighting the detailed venation pattern. **(C)** The root system, illustrating its typical cylindrical structure. **(D)** Overall habit of the plant, showing its characteristic branching pattern and foliage arrangement.

#### 3.1.2 Transverse Section (T.S) of leaf

T.S of leaf shows dorsiventral structure. The outer walls of the epidermal cells show the position of crystalline granules of calcium oxalate beneath the thick cuticle. Numerous multicellular glandular trichomes are present on both surfaces of the leaf. Mesophyll is distinguished into one layer of palisade cells and 2 to 4 layers of loosely arranged parenchyma. Anomocytic stomata present on both surfaces. Mid rib of leaf shows endodermis indistinct, pericycle 1-2 layered, thick-walled often containing scattered isolated fibres, stele consisting of many small vascular bundles often joined together in a ring and many big vascular bundles scattered in the ground tissue, intra fascicular cambium present. (Figure 2 (A))

**Figure 2.**
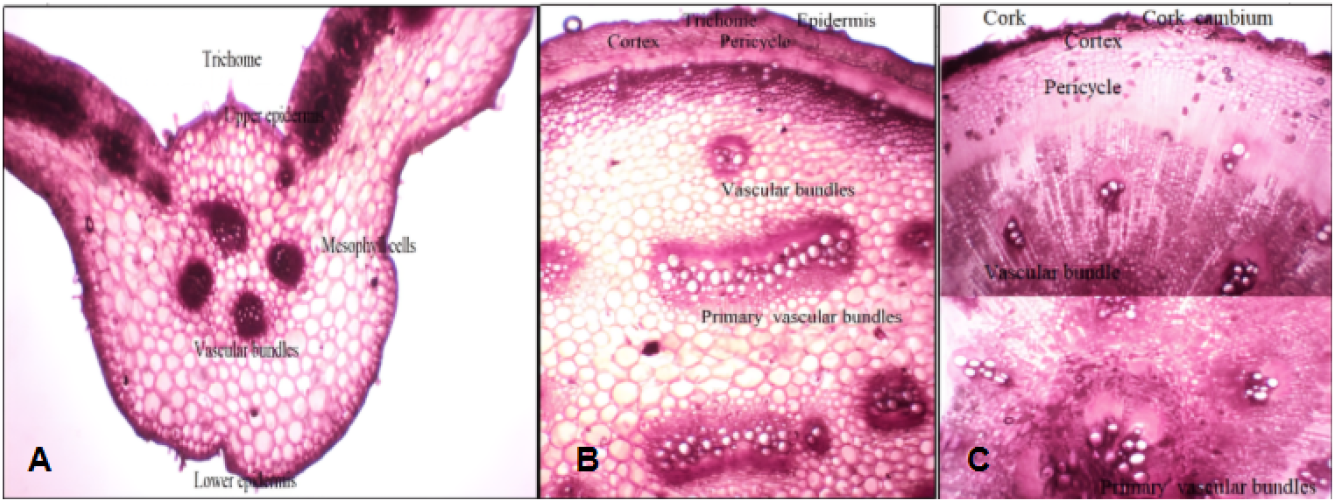
Anatomical features of *Boerhavia diffusa* in transverse section. (A) Leaf anatomy, (B) Stem anatomy, (C) Root anatomy

#### 3.1.3 Transverse section of Stem

The TS of the stem shows an epidermal layer bearing multi cellular, uniseriate glandular trichomes. The cortex comprised 1-2 layers of parenchyma cells, with an indistinct endodermis. The pericycle was 1-2 layers thick-walled, often containing scattered isolated fibres. The stele exhibited numerous small vascular bundles arranged in a ring and many big vascular bundles scattered in the ground tissue. Intrafascicular cambium was present. (Figure 2 (B))

#### 3.1.4 Transverse section of Root

The T.S of the matured root showed an outer cork layer composed of thin-walled tangentially elongated cells, with brown walls in the outer layers. The cork cambium consisted of 1-2 layers of thin walled cells. The secondary cortex comprised 2-3 layers of parenchymatous cells followed by a cortex composed of 5-12 layers of thin-walled, oval to polygonal cells. Below the cortex, several concentric bands of xylem tissue alternate with a wide zone of parenchymatous tissue. The number of xylem bands varied according to thickness of root and composed of vessels, tracheids and fibres. Vessels were predominantly arranged in radial groups of 2-8, exhibiting simple pits and reticulate thickening. Tracheids were small, thick-walled with simple pits. The central regions contained primary vascular bundles. Numerous calcium oxalate raphides, in single or in groups, were observed in the cortical region and parenchymatous tissue in between xylem tissue. Starch grains, both simple and compound, having 2-4 components were abundant throughout the cortical cells. (Figure 2 (C))

### 3.2 Simulated *in-vitro* digestion of PC samples found to release more iron compared to normal water extract

The samples’ iron content was measured via Inductively Coupled Plasma Mass Spectrometry (ICP-MS) to evaluate the differences in iron availability among various samples. The samples included blank (untreated), raw materials (leaf, whole plant and root), water extract of PC, and *in vitro* digested PC.

The iron concentration of raw materials varied across plant parts, with the root showing the highest content (1458.77 ppm), followed by the whole plant (1021.68 ppm) and leaves (290.57 ppm). Remarkably, after undergoing the *Ayolepam* process, all three preparations exhibited a several-fold increase in iron content, with leaves showing the highest (10504.79 ppm), followed by the whole plant (7353.95 ppm) and roots (5568.22 ppm).The findings are summarized in Table - 1.

**Table 1.**
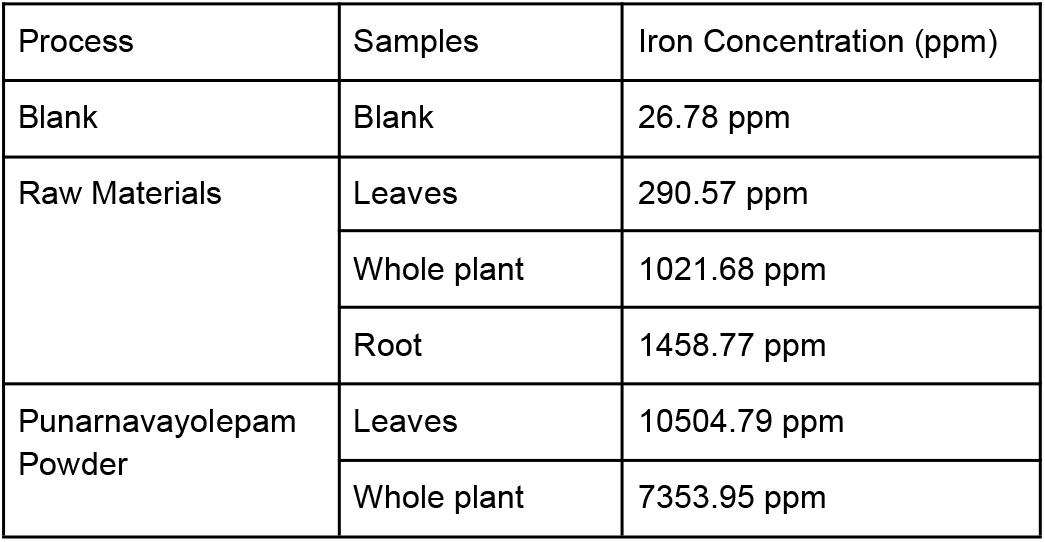
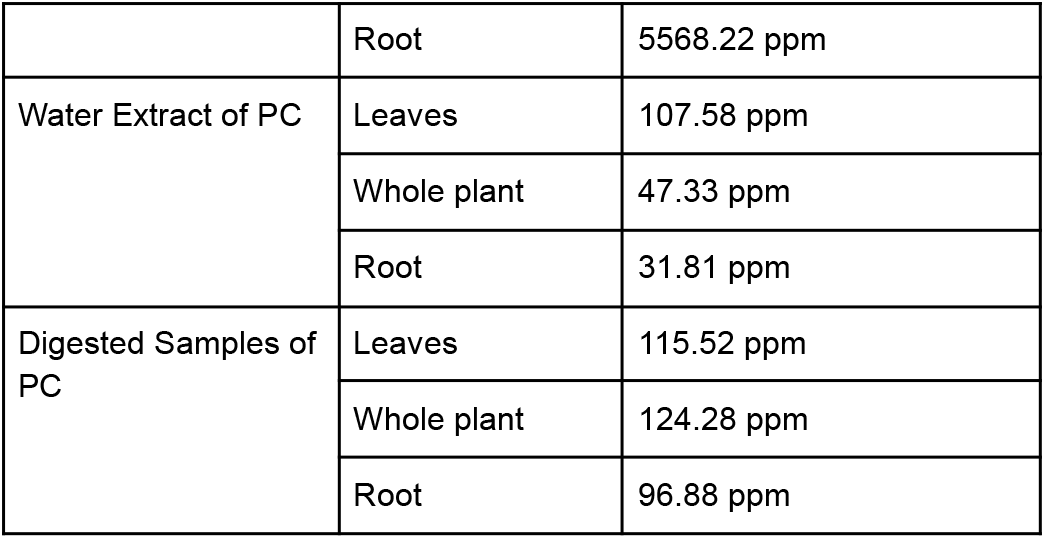
Iron concentration in *Boerhavia diffusa* materials and formulations. The table presents the iron concentration, measured in parts per million (ppm), across various samples. These include raw plant materials (*Boerhavia diffusa* leaf, whole plant, and root), Punarnavayolepam powder preparations, their corresponding water extracts, and extracts subjected to simulated in-vitro digestion. A key finding is that the simulated digestion of Punarnavayolepam (PC) samples resulted in significantly higher levels of released iron compared to the untreated water extracts. Blank values were included as controls for the analysis.

In order to understand the iron release from the PC, the samples were subjected to a simulated in vitro digestion and the digests were subjected to ICP-MS analysis. Alongside, a water extract of each sample, with the same amount, was prepared to compare the effect of iron release upon digestion. The iron concentration in the digested samples, regardless of the parts of *B. diffusa*, showed a statistically significant increase in iron compared to the water extract. The digests of whole-plant showed 124.28 ppm, while the leaves and roots showed 115.52 and 96.88 ppm respectively. At the same time, the water extracts of whole-plant, leaves and roots showed 47.33, 107.58 and 31.81ppm of iron release respectively (Table - 1).

### 3.5 PC digests showed improved bioavailability of iron in Caco-2 cells

Treatment of Caco-2 cells with PC digests, with or without FeSO_4_, showed a concentration dependent increase in intracellular iron concentration as quantified by ferrozine assay. Cells treated with 150µg/mL of FeSO_4_ showed a mean intracellular iron concentration of 22.59 ± 3.99 µg/mg of protein in one group (Fig - 3 A,C,E) and 19.27 ± 11.41 µg/mg of protein in the second group (Fig - 3 B,D,F). Treatment with all three samples of PC digests showed an increase in intracellular iron compared to the FeSO_4_ control. Among the preparations, the whole-plant extract demonstrated the highest intracellular iron concentration at 30% (122.72 ± 24.32 µg/mg of protein), followed by the leaf-based Ayolepam extract (104.98 ± 30.29 µg/mg of protein) and the root extract (73.18 ± 19.50 µg/mg of protein). Cells treated with samples alone (without added FeSO_4_) showed similar trend but with much lower amount of intracellular iron viz. 57.95±9.70 µg/mg of protein, 81.67±4.54 µg/mg of protein and 39.68±11.68 µg/mg of protein respectively for *B. diffusa* leaves, whole plant and roots. The lower concentrations of the sample showed a proportionate reduction of the intracellular iron concentration showing a linear relationship. All treatments with FeSO_4_ showed a significance in one-way ANOVA calculation whereas root digest treated without FeSO_4_ showed marginal increase in intracellular iron with no significance in one-way ANOVA analysis. Cells treated with 500µg/mL of ascorbic acid were used as a positive control for the experiment and it showed a significant increase of more than 2 fold iron absorption compared to the FeSO_4_ control. The results clearly indicate the iron bioavailability enhancing property of the preparation.

**Figure 3.**
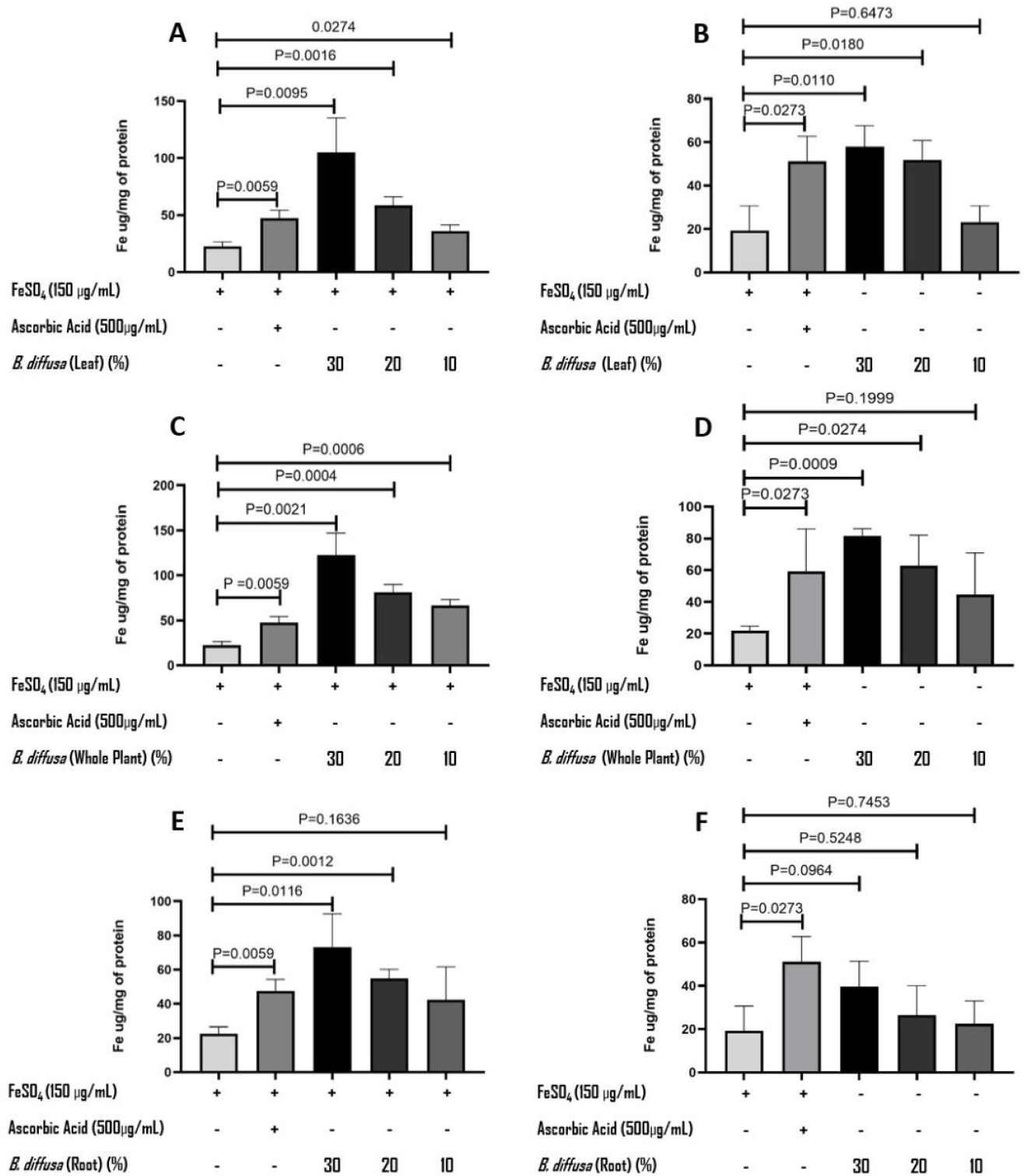
Effect of *Boerhavia diffusa* extracts on cellular iron uptake in CaCO_2_cells. CaCO_2_cells were treated with FeSO_4_ (150 µg/mL) and ascorbic acid (500 µg/mL) in the presence or absence of *B. diffusa* extracts at 10%, 20%, and 30% concentrations derived from (A, B) leaf, (C, D) whole plant, and (E, F) root. Cellular iron uptake was quantified and normalized to total protein, expressed as Fe µg/mg protein. Results are presented as mean ± SD from independent replicates. Statistical analysis was performed using one-way ANOVA followed by post-hoc multiple comparisons; significant p values are indicated.

## Discussion

Iron deficiency anemia is one of the major public health problems in developing countries. India, the country with the world’s largest adolescent population, has 67.1% of children and 59.1% of adolescent girls suffering from anemia according to the 5th National Family Health Survey [28]. The global scenario is not very different and the statistics show around 1.62 billion people affected with anemia [29]. While there are several strategies adopted across the globe for tackling IDA, the net result is not very encouraging as evidenced by the statistics. The major drawbacks associated with the current management strategies are poor bioavailability and assimilation of iron from diet and therapeutic interventions as well as low patient compliance of oral iron supplements [30]. To overcome these challenges, innovative pharmaceutic strategies can be adopted to develop interventions/supplements with better bioavailability, bioassimilation and patient compliance. In this direction, our study presents a unique pharmaceutical preparation called *Punarnavayolepam choornam* (PC), and its potential applications in IDA management.

The uniqueness of the PC is its pharmaceutics. It is a preparation of “*Ashta Vaidya*” school of *Ayurveda* from South India, where a single plant (*Boerhavia diffusa* - Punarnava) is processed on iron trays to entrap iron into the formulation. While the classical preparation uses the roots of *B. diffusa*, the current study explores the leaf and whole plant along as well. The pharmaceutical process showed a marked increase in iron concentration in PC prepared from leaf, whole-plant, and root samples following the *Ayolepam* process. This increase in the amount of iron in the PC preparation indicates the intrinsic iron-binding potential of the *B. diffusa* plant. It is quite interesting to note that in the raw state, higher amounts of iron was found in root, followed by the whole plant and leaves, whereas PC preparation following Ayolepam procedure showed the reverse order with leave preparation having highest amount of iron followed by whole plant and roots. More studies need to be carried out to understand the chemistry underlying this phenomenon of differential iron binding ability of different parts of a particular plant. Nevertheless, the fact that the root contains higher amounts of iron probably justify the rationale of selecting *B. diffusa* roots for PC preparation in the *Ashta Vaidya* school of pharmaceutics.

One of the major challenges in IDA management is the presence of anti-nutritive factors in dietary ingredients that reduce the iron bioavailability. These metabolites are known to sequester iron during gastro-intestinal digestion and significantly reduce the release of iron for absorption by intestinal cells [31]. In order to understand the dynamics of PC during digestion, a simulated *in-vitro* digestion model was employed in this study. The prepared PC (with leaf, root and whole plant) were subjected to simulated in-vitro digestion and the amount of iron released into the soluble part of the digest was measured. This was compared with a sham digestion with an equivalent amount of distilled water. This step is critical, as plant-derived iron is typically complexed with phytates, fibres, proteins, and polyphenols, rendering it insoluble and poorly available for absorption [32]. Here again the observation was quite interesting that the leaf based PC preparation showed a similar amount of iron release with water and complete digestion, whereas both whole plant and root showed a significant increase in iron release after digestion, as compared to sham (water extract). This result suggests that *B. diffusa* is a plant that not only increases the absorption of iron from the source (here for PC preparation, the iron tray) but it also releases iron after digestion and makes it ready for bioavailability. This makes PC an ideal pharmaceutical preparation for IDA management.

The classical preparation of PC uses *B. diffusa* roots, but our study observed the iron content is more in leaf and whole-plant based preparations. It is intuitive to assume that replacing *B. diffusa* roots with its leaf or whole-plant may have impart better pharmacological properties to PC with respect to IDA management. However, it is important to note the possible anti-nutritive factors (ANFs) present in different parts of *B. diffusa* that may hamper the bioavailability and assimilation of iron. The aerial part of *B. diffusa* contains tannins, phenols, and phytic acid that are shown to inhibit iron absorption, especially non-heme iron, rendering it unavailable for assimilation [33]. Studies indicate that when compared to leaves and stem, the *B. diffusa* Roots exhibited the lowest concentration of these anti nutrition factors with tannins (0.45%), saponins (0.77%), and phenols (0.09%) [34] Similarly, *B. diffusa* leaves also contain high levels of minerals, particularly calcium (667 mg/100g) [35]. Such elevated Calcium levels are also shown to potentially compete with iron and inhibit its absorption [36].

Additionally, it is very important to note that *B. diffusa* roots contains some of the beneficial phytoconstituents, such as rotenoids, flavonoids, flavonoid glycosides, xanthones, purine nucleosides, lignans, ecdysteroids, and steroids [37]. Many of these compounds have been studied in facilitating non-heme iron absorption and mobilization through antioxidant activity, chelation, or interaction with cellular transport mechanisms. Studies indicate that the root of *B. diffusa* had the maximum antioxidant activity compared to the leaf and stem samples [38]. Our iron bioavailability study using Caco-2 cells demonstrates that treatment with digests from all three samples of the PC preparation effectively enhanced intracellular iron uptake compared to the FeSO4 control, suggesting a superior bioavailability. While this enhanced uptake is beneficial, it is crucial to maintain moderate iron supplementation. It is also a known fact that excess amount of iron in the body impacts the metabolic homeostasis negatively by damaging various organs (hemochromatosis) and therefore it is important to have moderate iron supplementation where the body absorbs maximum and reduces the wastage [39]. Although we do not have any clear justification for the Ayurvedic perspective of opting *B. diffusa* roots over leaf or whole-plant, ignoring the possible sustainable harvesting challenges, our analysis provides an acceptable explanation for the use of *B. diffusa* roots for PC preparation. The root based preparations are clinically proven and the mode of action studies conducted are also supporting the clinical claims. However, more detailed studies can probably shed more light to establish the efficacy and suitability of leaf or whole-plant based PC over root from both pharmacology and pharmaceutics point-of-view.

The Recommended Dietary Allowance (RDA) of iron by US National Institutes of Health is set at 8 mg/day for healthy adult men and 18 mg/day for premenopausal women [40]. The ICMR-National Institute of Nutrition (NIN) recommends iron for adult men as 19 mg/day and non-pregnant, non-lactating women as 29 mg/day which is higher than the international standards as the iron considering the lower bioavailability of non-heme iron in the typical Indian diet [41]. Our PC preparations can provide support in meeting the RDA of iron for populations at risk of iron deficiency anemia. PC root preparation containing 5.57 mg of iron per gram, administered at a daily dose of 10 g, provides an adequate amount to meet the requirements. Also for treating IDA, Ashtavaidya’s practice of giving buttermilk as the adjuvant for PC again supports in improving the bioavailability of iron. The fermented buttermilk can reduce phytates and other anti-nutritional factors, which otherwise can limit iron absorption and can enhance the micronutrient availability including iron and zinc [42].

The milk fat globule membrane components (such as sphingolipids and gangliosides) that modulates iron transport, accompanied with antioxidant and reducing activity from high sulfhydryl content which binds and retain iron in the absorbable ferrous (Fe^2+^) state. Furthermore, the presence of phosphoserine residues in milk proteins (especially caseins) helps in strong iron binding: fermentation again hydrolyzes these proteins, while the antioxidants protect iron from oxidation and lipid peroxidation, combinely boosting intestinal iron uptake [43] [44].

## Conclusion

Collectively, our results present a novel pharmaceutical product-design from a traditional school of Ayurveda and these results demonstrate its iron-mobilizing and absorption-enhancing potential through in-vitro model systems. Our study reaffirms the traditional use of *Punarnava* as a Rasayana herb in Ayurveda and highlights its potential in iron deficiency anemia management. Given the global burden of iron deficiency anemia, further translational studies, including in vivo studies and human intervention trials along with the adjuvant, are required to substantiate the efficacy of *Punarnavayolepa Choornam* (PC) as functional nutritional interventions for improving iron status.

## Supporting information

https://www.compress2go.com/result#j=b97d313f-0f8f-4627-ab6a-8b0359f7b846

## Acknowledgement

The authors gratefully acknowledge the financial support provided by Vaidyaratnam Ousadhasala,Thaikkattussery, Thrissur, Kerala. The authors CNVP and SKK acknowledge the financial support received from the Rural India Support Trust (RIST) through TDU. The authors also acknowledge the financial support for a Research Fellow from The University of Trans-Disciplinary Health Sciences and Technology (TDU), Bangalore.

## Author Contributions

SKB, CNVP, SKK, AGR and JGN conceptualized the work; experiments were performed by JGN, RTS, DM, ABS, PIC, CNVP, AGR and VW; data validation was done by SKB, CNVP, YNM, RVM, AA, SKK, JGN, ABS and AGR.; resources for the experiments were arranged by CNVP, AGR, YNM, RVM and AA.; Data curation and original manuscript writing was done by JGN, RTS and CNVP and review of manuscript was done by CNVP, YNM, RVM, AA, SKK, SKB, JGN and AGR. All authors have read and agreed to the published version of the manuscript.

