## Supplementary material for "“*Punarnavayolepa Choornam*” in Iron Deficiency Anaemia management: Pharmaceutic insights and biological activity": https://www.compress2go.com/result#j=b97d313f-0f8f-4627-ab6a-8b0359f7b846: Supplementary Files.pdf

**1.** Powder microscopy

**Table 1**:Comparative Powder Microscopic characters of raw leaf and processed leaf(Ayolepa leaf) samples are summarized below

| No | Microscopic features | Raw leaf (A) | Processed leaf (B) |
|---|---|---|---|
| 1. | Starch grains | Present | Present |
| 2. | Fragment of mesophyll cells with contents | Present | Present |
| 3. | Prismatic crystal | Present | Present |
| 4. | Acicular crystals | Present | Comparatively lesser |
| 5. | Oil globule | Present | Present |
| 6. | Trichome | Present | Present |
| 7. | Dark coloured cellular masses | | Numerous |
| 8. | Stomata | Present | Present |

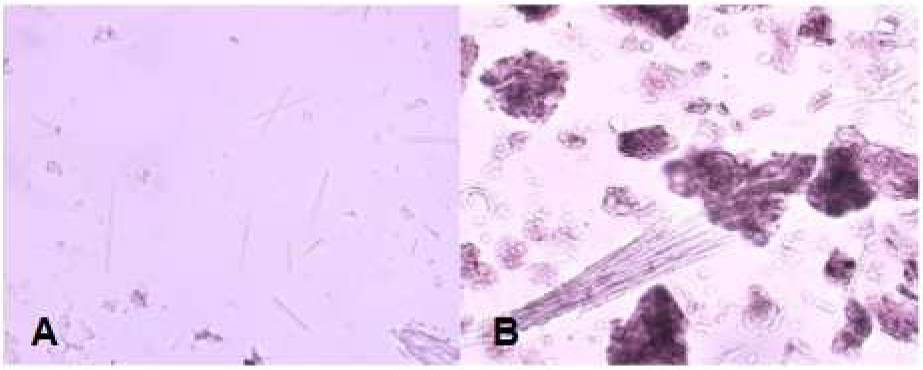


**Figure 1:** Comparative Powder Microscopic characters of raw leaf and processed leaf(Ayolepa leaf)

**Table 2:** Comparative Powder Microscopic characters of raw root and processed root(Ayolepa root) samples are summarized below.

| No | Microscopic features | Raw root (C) | Processed root (D) |
|---|---|---|---|
| 1. | Starch grains | Numerous | Numerous |
| 2. | Acicular crystals | Numerous | Comparatively lesser |
| 3. | Prismatic crystals | Present | Present |
| 4. | Vessel fragments | Present | Present |
| 5. | Fibres | Present | Present |
| 6. | Tracheids | Present | Present |
| 7. | Oil globule | Present | Present |

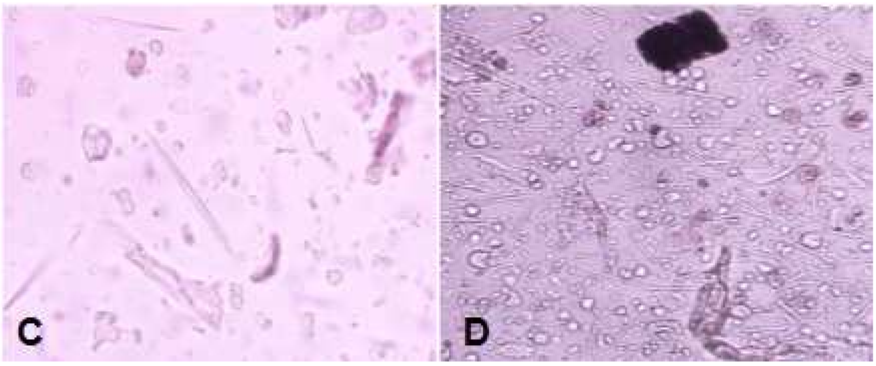


**Figure 2:** Comparative Powder Microscopic characters of raw root and processed root(Ayolepa root)

**Table 3:** Comparative Powder Microscopic characters of raw whole plant and processed whole plant (Ayolepa whole plant ) samples are summarized below.

| No | Microscopic features | Raw whole plant (E) | Processed whole plant (F) |
|---|---|---|---|
| 1. | Starch grains | Numerous | Present |
| 2. | Acicular crystals | Numerous | Numerous |
| 3. | Prismatic crystals | Present | Present |
| 4. | Vessel fragments | Present | Present |
| 5. | Fibres | Present | Present |
| 6. | Tracheids | Present | Present |
| 7. | Oil globule | Present | Present |
| 8. | Fragment of mesophyll cells with cellular contents | Present | Present |
| 9. | Trichome | Present | Present |

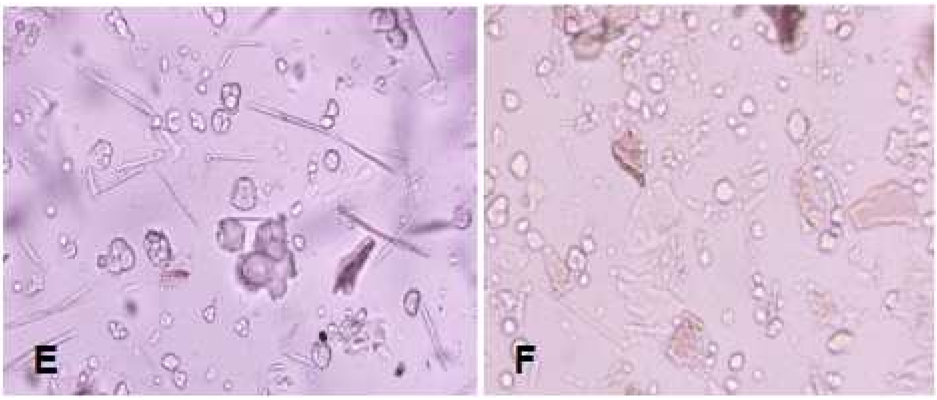


**Figure 3:** Comparative Powder Microscopic characters of raw whole plant and processed whole plant

Boerhaavia diffusa powder microscopy shows that acicular (needle-shaped) calcium oxalate crystals are present in all plant parts and are a distinguishing feature. However, the processed (Ayolepa) leaf and root samples show an apparent reduction in the quantity of these crystals. Furthermore, there are notable microstructural differences between the raw and processed forms of the plant due to the relative abundance of starch grains in the root parts.

**2. Physicochemical Analysis**
The physicochemical examination of the formulation was conducted to determine its moisture content, total ash value, and carbohydrate composition, with the results provided in table 4.

Table 4: Moisture content, total ash value, and carbohydrate composition of different samples.

| SL No. | Sample Details | LOD % | Total Ash % | Total Carbohydrate % |
|---|---|---|---|---|
| 1 | Raw leaves | 4.61% | 17.53% | 77.86% |
| 2 | Leaves (Paste smeared) | 2.62% | 18.67% | 78.71% |
| 4 | Raw whole plant | 3.65% | 9.65% | 86.70% |
| 5 | Whole plant (Paste smeared) | 1.85% | 12.21% | 85.94% |
| 7 | Raw Root | 3.32% | 6.92% | 89.76% |
| 8 | Root (Paste smeared) | 2.57% | 8.70% | 88.73% |

**3. Nutritional Profiling**

Table 5: The table represents the nutrition profile of different samples based on the different parts of the Punarnava plant. Sodium, potassium and calcium are calculated in ppm level.

| No. | Sample details | Sodium | Potassium | Calcium | Calorific value |
|---|---|---|---|---|---|
| 1 | Raw leaves | 0.36 ppm | 5.44 ppm | 1.80 ppm | 311.44 |
| 2 | Leaves (Paste smeared) | 0.31 ppm | 4.51 ppm | 1.81 ppm | 314.84 |
| 4 | Raw whole plant | 0.35 ppm | 2.78 ppm | 1.17 ppm | 346.8 |
| 5 | Whole plant (Paste smeared) | 0.33 ppm | 2.74 ppm | 1.66 ppm | 343.76 |
| 7 | Raw Root | 0.38 ppm | 0.98 ppm | 1.40 ppm | 359.04 |
| 8 | Root (Paste smeared) | 0.33 ppm | 1.14 ppm | 1.77 ppm | 354.92 |
